# Activation of prodynorphin neurons in the dorsomedial hypothalamus inhibits food intake and promotes positive valence

**DOI:** 10.1101/2020.07.02.183780

**Authors:** Daigo Imoto, Izumi Yamamoto, Hirokazu Matsunaga, Toya Yonekura, Ming-Liang Lee, Kan X Kato, Takeshi Yamasaki, Ken-ichi Otsuguro, Motohiro Horiuchi, Norifumi Iijima, Kazuhiro Kimura, Chitoku Toda

**Affiliations:** Laboratory of Biochemistry, Graduate School of Veterinary Medicine, Hokkaido University, Sapporo, Hokkaido, 060-0818, Japan; Laboratory of Animal Experiment, Institute for Genetic Medicine, Hokkaido University, Sapporo 060-0815, Japan; Laboratory of Pharmacology, Graduate School of Veterinary Medicine, Hokkaido University, Sapporo 060-0818, Japan; Laboratory of Veterinary Hygiene, Graduate School of Veterinary Medicine, Hokkaido University, Sapporo 060-0818, Japan; National Institutes of Biomedical Innovation, Health and Nutrition, Ibaraki, Osaka, 567-0085, Japan; Immunology Frontier Research Center, Osaka University, Suita, Osaka, 565-0871, Japan

## Abstract

The regulation of food intake is one of the major research areas in the study of metabolic syndromes such as obesity. Gene targeting studies have clarified the roles of hypothalamic neurons in feeding behaviour. However, our understanding of neural function under physiological conditions is still limited. Immediate early genes, such as activity-regulated cytoskeleton-associated protein (Arc/Arg3.1), are useful markers of neuronal activity. Here, we investigated the role of Arc/Arg3.1 gene-expressing neurons in the hypothalamus after refeeding using the targeted recombination in active populations method. We identified refeeding-responsive prodynorphin/cholecystokinin neurons in the dorsomedial hypothalamus that project to the paraventricular hypothalamic nucleus. Chemogenetic activation of these neurons decreased food intake and promoted positive valence. Our findings provide insight into the role of newly identified hedonic neurons in the process of feeding-induced satiety.

## Introduction

An imbalance between food intake and energy expenditure leads to various health problems, such as obesity. Food intake is regulated by homeostatic controls of appetite that induce hunger (feeding), satiation (suppression of feeding), and satiety (post-meal termination of hunger)^1,2^. Recent studies have identified neurons and neuronal circuits that regulate food intake^3–6^. For example, the arcuate nucleus of the hypothalamus (ARC) contains various cell types, and receives endocrine and exogenous signals that regulate feeding behaviours. Ghrelin secreted from the stomach activates neurons that release neuropeptide Y (NPY) and agouti-related protein (AgRP) in the ARC, resulting in increased food intake^7^. In contrast, the activation of proopiomelanocortin (POMC)-releasing neurons and vesicular glutamate transporter 2 (VGLUT2)-expressing neurons in the ARC decreases food intake^8–10^. The paraventricular nucleus of the hypothalamus (PVH) contains two subnuclei—the ventral parvocellular and dorsal magnocellular regions—that have distinct neuronal projections. A subset of PVH neurons, expressing pituitary adenylate cyclase-activating polypeptide (PACAP) and thyrotropin-releasing hormone (TRH), activates NPY/AgRP neurons via glutamatergic neurotransmission^11^. In addition, NPY/AgRP neurons suppress other types of PVH neurons to promote satiety^11^. Although the dorsomedial hypothalamus (DMH) is less well-studied than the ARC and PVH, NPY neurons and cholinergic neurons in this region are involved in the regulation of food intake^12,13^.

Most studies on the neural regulation of feeding have used cell-type-specific knockout performed with the Cre recombinase (Cre)/loxP system. While it is a powerful method for investigating the role of specific neurons and neuropeptides, it has methodological limitations. For example, the permanent deletion of a gene has a long-term effect on neurophysiology, making it unsuitable for investigating short-term changes such as appetite. Furthermore, gene deletions during the developmental stage may affect neuronal projections and brain structures. Therefore, the role of these neuronal circuits in the regulation of food intake under normal physiological conditions remains unclear.

Recently, Guenthner et al. developed a new approach for labelling active neurons *in vivo*, called the targeted recombination in active populations (TRAP)^14^ method. In this method, the promoter of the cFos or activity-regulated cytoskeleton-associated protein (Arc/Arg3.1) gene, which are members of the immediate early gene family that are used as markers of neuronal activity, is used to express tamoxifen-dependent Cre. Tamoxifen injection induces the expression of a fluorescent protein in active neurons expressing cFos or Arc/Arg3.1. We selected Arc/Arg3.1 because it is a neuron-specific gene, while cFos is not.

In this study, we investigated the role of active hypothalamic neurons during the refeeding period using the TRAP method^14^. We observed an increase in TRAP-labelled neurons in the DMH, 1 h after refeeding. The chemogenetic activation of DMH neurons by excitatory designer receptors exclusively activated by designer drugs (DREADDs) decreased food intake, suggesting that these neurons are involved in the regulation of feeding behaviour in mice. In addition, the activation of these neurons produced conditioned place preference (CPP), suggesting that they are involved in producing the reward effect (positive valence) during feeding. The RNA sequencing study revealed that the labelled neurons in the DMH expressed an opioid polypeptide, prodynorphin (pdyn), as well as cholecystokinin (CCK). Taken together, our findings reveal a novel type of neuron in the DMH that regulates the appetite satiety process and promotes positive valence in mice. This neuronal circuit could be a potential therapeutic target for obesity and binge eating disorder.

## Materials and methods

### Animal ethics and husbandry

All animal experiments were approved by the Animal Care and Use Committee of Hokkaido University, and were performed according to the institutional guidelines. Arg3.1-Cre/ER^T2^ and Ai14 mice were purchased from The Jackson Laboratory (Bar Harbor, ME, USA). In this study, Arg3.1-Cre/ER^T2^ and Arg3.1-Cre/ER^T2^;Ai14 mice (8–20 weeks of age) were used. Arg3.1-Cre/ER^T2^ mice were crossed to Ai14 mice to obtain Arg3.1-Cre/ER^T2^;Ai14 mice. All mice were kept at 22 ± 4 °C under a 12/12-h light/dark cycle (light phase: 7:00–19:00) and given *ad libitum* food and water access. Mice were fed regular chow diet (RCD) from Nosan Corporation (Yokohama, Japan).

### Targeted recombination in active populations (TRAP)

Male Arg3.1-Cre/ER^T2^;Ai14 mice were fasted for 16 h and fed RCD *ad libitum*. The mice received 4-hydroxy tamoxifen (4-OHT) (10 mg/kg; Sigma-Aldrich, St. Louis, MO, USA) intraperitoneal (i.p.) injection at 0, 0.5, 1 or 2 h after refeeding to label neurons expressing the Arc/Arg3.1 gene.

### Brain sectioning and immunohistochemistry

Mice were sacrificed using CO_2_ asphyxiation and perfused with heparinized saline. Brain samples were harvested and incubated in 4% paraformaldehyde (PFA) overnight. Brain sections (50 μm) containing the whole hypothalamus were collected. Floating sections were incubated with rabbit anti-cFos antibody (1:200; Santa Cruz Biotechnology, Denton, TX, USA), rabbit anti-GFP antibody (1:1,000; Frontier Institute, Ishikari, Japan), mouse anti-cFos antibody (1:1,000; Santa Cruz Biotechnology) or rabbit anti-dynA antibody (1:200; Phoenix Pharmaceuticals, Burlingame, CA, USA) in staining solution (0.1 M phosphate buffer [PB]) overnight at room temperature. After rinsing with PB, sections were incubated with Alexa 488 anti-rabbit (IgG) (1:250; Life Technologies, Carlsbad, CA, USA) or Alexa 594 anti-mouse (IgG) (1:250; Cell Signaling Technologies, Danvers, MA, USA) for 2 h at room temperature. Brain sections stained with guinea pig anti-tdTomato antibody (1:1,000; Frontier Institute) were incubated with biotin-labelled anti-guinea pig IgG (1:250; Invitrogen, Carlsbad, CA, USA) overnight. After rinsing with PB, sections were incubated with Alexa 594 streptavidin (1:2,000; Life Science Technologies, Eugene, OR, USA) for 1 h at room temperature. Sections were then rinsed with PB and mounted on glass slides with Vectashield (Vector Laboratories, Burlingame, CA, USA). Fluorescent signals for EGFP and mCherry in Figures 2B (EGFP: rabbit anti-GFP; mCherry: guinea pig anti-tdTomato) and 4B–D (EGFP: rabbit anti-GFP) were enhanced by immunohistochemistry for visualization. Quantification of fluorescent signals was carried out using ImageJ^15^. Image colours used in Figure 2B were adjusted; EGFP (green) to red and cFos (red) to green using ImageJ for comparison.

**Figure 1.**
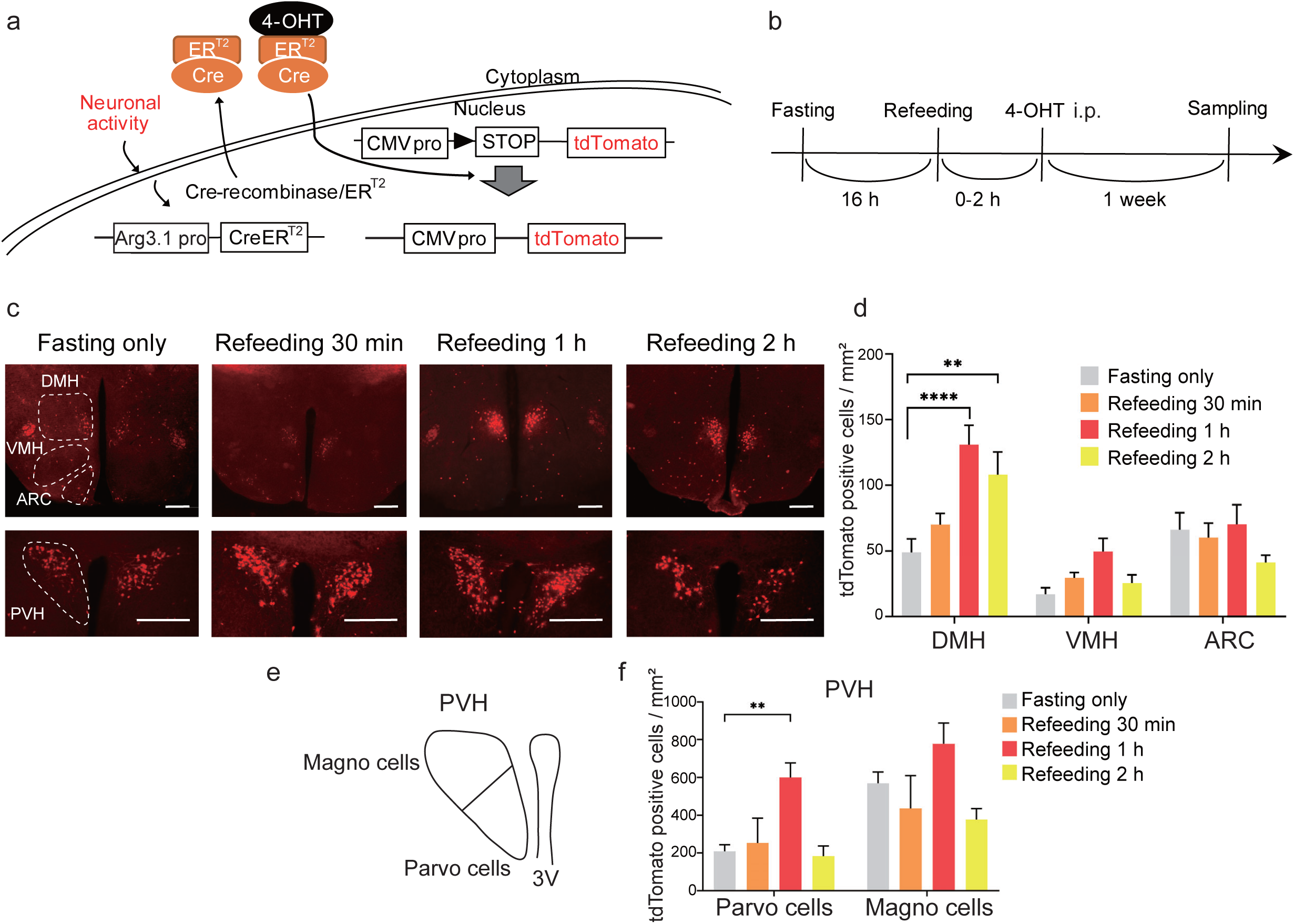
Fasting-refeeding activates neurons in the DMH. **a** Genetic strategy for visualizing activated neurons using the TRAP method. Neuronal activity promotes expression of 4-hydroxy tamoxifen (4-OHT)-dependent Cre recombinase (Arg3.1-Cre/ER^T2^). The recombination of the reporter gene occurs in the presence of 4-OHT, allowing the expression of the tdTomato reporter gene for fluorescence visualization. **b** Experimental scheme. Male Arg3.1-Cre/ER^T2^;Ai14 mice were fasted overnight (16 h) and re-fed regular chow diet (RCD). 4-OHT intraperitoneal (i.p.) injection was given at 0, 0.5, 1 or 2 h after refeeding. Samples were collected 1 week after the injection. **c** Representative fluorescence images of TRAP-labelled tdTomato-positive neurons in the DMH, VMH, ARC and PVH at 0, 0.5, 1 or 2 h after refeeding. Scale bar: 300 μm. **d** Quantification of TRAP-labelled neurons in the DMH (*n* = 4–6), VMH (*n* = 4 or 5) and ARC (*n* = 3) at 0, 0.5, 1 or 2 h after refeeding. **e** Schematic showing the parvocellular and magnocellular regions of the PVH. **f** Quantification of TRAP-labelled neurons in the parvocellular and magnocellular regions of the PVH (*n* = 3– 5). All data are shown as mean ± SEM. Two-way ANOVA, Tukey post hoc test. \*\**P* < 0.01; \*\*\*\**P* < 0.0001.

**Figure 2.**
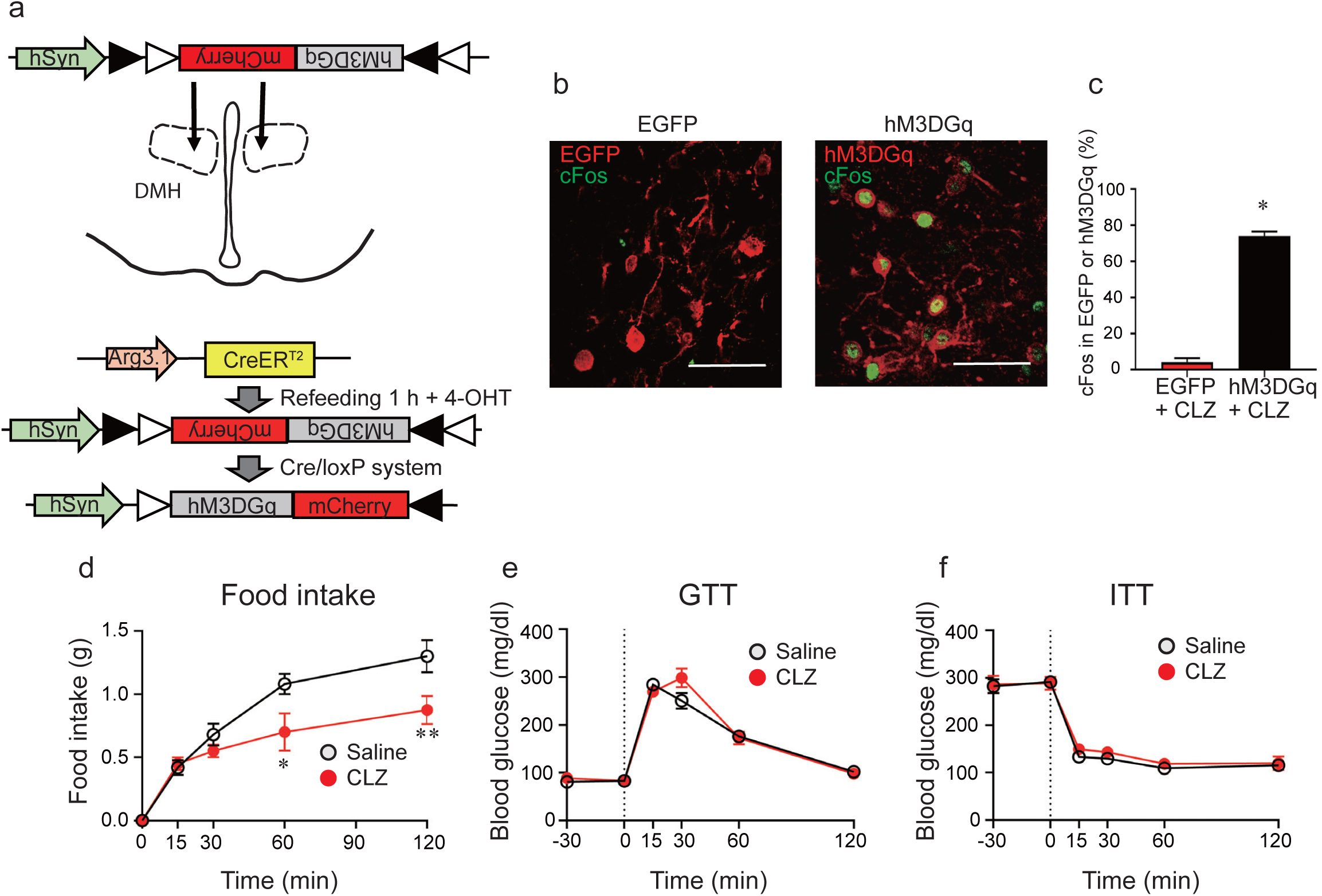
Chemogenetic activation of labelled DMH neurons decreases food intake without affecting glucose metabolism. **a** The AAV vector construct used for this experiment. Male Arg3.1-CreER^T2^ mice received bilateral AAV2-hSyn-DIO-hM3DGq-mCherry virus injection into the DMH. After recovery, the mice were fasted overnight and re-fed RCD. 1 h after refeeding, mice were given 4-OHT injection (i.p.) to induce Cre recombination. **b** Co-localization of cFos (green) and EGFP (red) or hM3DGq (red) after CLZ injection. Scale bar: 50 μm. **c** Quantification of the colocalization of cFos and EGFP (*n* = 3) or hM3DGq (*n* = 5) (mean ± SEM). **d** Food intake (0–120 min) after i.p. injection (−30 min) of CLZ (*n* = 4) or saline (*n* = 5). **e** Glucose tolerance test (GTT) (0–120 min) after i.p. injection (−30 min) of CLZ (*n* = 7) or saline (*n* = 7). **f** Insulin tolerance test (ITT) (0–120 min) after i.p. injection (−30 min) of CLZ (*n* = 5) or saline (*n* = 7). Each point represents mean ± SEM. Two-way ANOVA, Sidak post hoc test. \**P* < 0.05.

### Stereotaxic surgeries and adeno-associated virus (AAV) injection

Arg3.1-Cre/ER^T2^ mice (12–20-week-old) were anesthetized with a mixture of ketamine (100 mg/kg) and xylazine (10 mg/kg), and positioned on a stereotaxic instrument (Narishige, Tokyo, Japan). Mice were injected in each side or one side of the DMH with ∼0.5 µL AAV2-hSyn-DIO-hM3D(Gq)-mCherry (Addgene, Cambridge, USA), AAV2-hSyn-DIO-hM4D(Gi)-mCherry (Addgene), AAV8-eSyn-DIO-EGFP (Addgene) or AAV8-eSyn-DIO-eNpHR3.0-EGFP (Vector Biolabs, Malvern, PA, USA) using the following coordinates: AP: −1.85 mm, L: ± 0.3 mm, DV: −5.5 mm. Open wounds were sutured after viral injection. A 7–14-day recovery period was allowed before experiments were started. Mice were fasted overnight (16 h) and received 4-OHT (10 mg/kg) injection (i.p.) 1 h after refeeding to induce Cre recombination. To ensure adequate expression of proteins, experiments were carried out at least 7 days after the 4-OHT injection.

### Glucose and insulin tolerance test

Glucose and insulin tolerance tests were performed on fasted mice expressing DREADDs (hM3DGq or hM4DGi) to assess the role of refeeding-responsive DMH neurons. To activate DREADDs, clozapine (CLZ) solution (1 mg/kg; Toronto Research Chemicals, North York, ON, Canada) was injected (i.p.) 30 min before glucose or insulin tolerance tests. For the glucose tolerance test, animals were fasted for 16 h and injected with glucose (2 g/kg; i.p.). Blood glucose levels were measured with a handheld glucose meter (Nipro Free Style, Nipro, Osaka, Japan) before injection of CLZ (−30 min) and glucose (0 min), as well as 15, 30, 60 and 120 min after glucose injection. The insulin tolerance test was performed on *ad libitum*-fed mice. The mice were injected with 0.75 U/kg insulin (Novo Nordisk, Bagsværd, Denmark). Blood glucose was measured before injection of CLZ (−30 min) and insulin (0 min), as well as 15, 30, 60 and 120 min after insulin injection.

### Food intake measurement

Animals were placed in a new cage (single cage housing) the day before the experiment. Food was removed before the beginning of the dark phase, and the mice were fasted for 16 h. Animals received saline or CLZ (1 mg/kg) injection (i.p.) 30 min before refeeding. Food intake was measured at 0, 15, 30, 60 and 120 min.

### Conditioned place preference (CPP) test

The CPP apparatus consisted of compartments distinguished by a yellow striped floor (with stripes) and a white floor (no stripes) connected by an adjacent aisle. All experiments were carried out in the light phase. The protocol for the CPP test was as described by Kim et. al^16^. Test and conditioning times were measured with a stopwatch, and the video recording was analysed using ImageJ^15^ and MouBeAT^17^. On day 1, male Arg3.1-Cre/ER^T2^ mice expressing hM3DGq or EGFP were placed in the CPP apparatus and allowed to explore freely for 10 min without saline or CLZ injection to determine intrinsic place preference. Animals with intrinsic place preference index (PPI) > 50% were excluded from the analysis. PPI (%) was calculated using Equation 1.

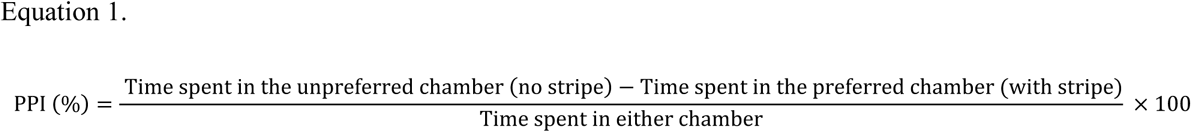

Conditioning took place on days 2–5, where the animals had access to only one chamber. On days 2 and 4, the mice received CLZ injection (i.p.) 40 min prior to the experiment and were placed in a chamber without stripes for 30 min. On days 3 and 5, the mice received saline injection (i.p.) 40 min prior to the experiment and were placed in a chamber with stripes for 30 min. On day 6, the mice freely explored the entire CPP apparatus for 10 min without saline or CLZ injections, and the time spent in each chamber was video-recorded. Normalized place preference was calculated using Equation 2.

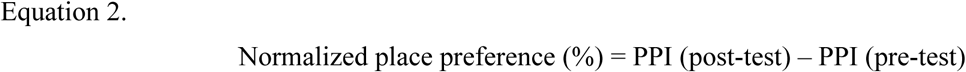

### Cell sorting for RNA sequencing

TRAP-labelled refeeding-responsive DMH neurons were collected by sorting tdTomato-positive neurons from the DMH of male Arg3.1-Cre/ER^T2^;Ai14 mice. Male Arg3.1-Cre/ER^T2^;Ai14 mice (8–12-week-old) were fasted overnight (16 h) and given 4-OHT injection 1 h after refeeding with RCD the next day. Sample collection was carried out 1 week after the injection.

We used a modification of a protocol reported previously for manually sorting tdTomato-positive and negative neurons^18^. Mice were sacrificed using CO_2_ asphyxiation, and the brain was placed in ice-cold cutting solution (220 mM sucrose, 2.5 mM KCl, 6 mM MgCl_2_, 1 mM CaCl_2_, 1.25 mM NaH_2_PO_4_, 10 mM glucose, 26 mM NaHCO_3_, bubbled thoroughly with 95% O_2_/5% CO_2_). A coronal brain slice (500 μm) containing the DMH was obtained using a vibratome (PELCO easiSlicer, Redding, CA, USA). The DMH was dissected and placed in a tube containing filtered artificial cerebrospinal fluid (ACSF; 105 mM NaCl, 2.5 mM KCl, 2 mM CaCl_2_, 1.3 mM MgSO_4_, 1.23 mM NaH_2_PO_4_, 24 mM NaHCO_3_, 20 mM HEPES, 2.5 mM glucose, 100 nM TTX, 20 μM DQNX, 50 μM AP-V, pH 7.4, bubbled thoroughly with 95% O_2_/5% CO_2_) with papain (0.3 U/ml), DNase (0.075 μg/ml) and BSA (3.75 μg/ml). The tube was incubated on a shaker (34 °C, 75 rpm) for 15 min. After incubation, the tissue was washed three times with papain-free filtered ACSF supplemented with FBS (1%). Then, 1 ml ACSF with FBS (1%) was added, and the samples were triturated successively with 600, 300 and 150 μm fire-polished Pasteur pipettes (10 times each). The tube was centrifuged (120 x g, 5 min, room temperature), and the supernatant was removed. The cell pellet was resuspended in 10 ml filtered ACSF containing FBS (1 %) and transferred to a 100 mm collagen type I-coated petri dish. The cells were allowed to settle onto the floor of the dish for 10 min, and tdTomato-positive and negative neurons were sorted separately under the fluorescence microscope using a cell aspirator attached to a micropipette (diameter: 30– 50 μm). The neurons were transferred to a clean 35 mm collagen type I-coated petri dish containing filtered ACSF with FBS (1%). This sorting process was repeated one more time, and cells were transferred to a 35 mm collagen type I-coated petri dish containing filtered PBS (bubbled thoroughly with 95% O_2_/5% CO_2_). The sorted neurons were transferred to a PCR tube (five neurons per tube) containing 1 μL of 10× reaction buffer (SMART-Seq v4 Ultra Low Input RNA Kit for Sequencing, Cat. No. 634888, Takara Bio, Kusatsu, Japan). Nuclease-free water was added to 11.5 μL, and the mixture was stored at −80 °C until library preparation.

### RNA sequencing

RNA sequence library preparation, sequencing, and mapping of gene expression were performed by DNAFORM (Yokohama, Kanagawa, Japan). Double-stranded cDNA libraries (RNA-seq libraries) were prepared using SMART-Seq v4 Ultra Low Input RNA Kit for Sequencing (Cat. No. 634888, Takara Bio) according to the manufacturer’s instructions. RNA-seq libraries were sequenced using paired-end reads (50-nt read 1 and 25-nt read 2) on a NextSeq 500 instrument (Illumina, San Diego, CA, USA). Obtained reads were mapped to the mouse GRCm38.p6 genome using STAR (version 2.7.3a). Reads on annotated genes were counted using featureCounts (version 1.6.4). FPKM values were calculated from mapped reads by normalizing to total counts and transcript. Prism 8 software (GraphPad) was used to generate heatmaps. Gene lists used for RNA sequencing analysis were obtained from previous reports^19,20^.

### Statistical analysis

All data are presented as the mean ± SEM, and *n* represents the number of animals. Statistical differences were evaluated with Student’s unpaired *t-*test (for two-group comparisons) or two-way ANOVA. Tukey or Sidak post-hoc tests (for multiple comparisons) were performed using Prism 8 software (GraphPad). Values of *P* < 0.05 were considered significant.

## Results

### Fasting-refeeding activates a subset of neurons in the compact part of the DMH

To identify neurons activated by refeeding, the TRAP method was used to label Arg3.1-expressing neurons. This method requires two transgenes; one expresses Cre recombinase fused with oestrogen receptor type 2 (Arg3.1-Cre/ER^T2^) from an activity-dependent Arc/Arg3.1 promoter, and the other allows the expression of a tdTomato reporter for fluorescence visualization (Ai14:loxP-stop-loxP-tdTomato). 4-OHT, an active form of tamoxifen, was injected to induce translocation of Cre into the nucleus to cause recombination of the reporter gene. Continuous expression of the fluorescent proteins is driven by the cytomegalovirus (CMV) promoter (Fig. 1a).

Male Arg3.1-Cre/ER^T2^;Ai14 mice were fasted overnight and re-fed RCD. 4-OHT was injected (i.p.) at 0, 0.5, 1 or 2 h after refeeding, and the brain samples were collected 1 week later (Fig. 1b). In the fasting only condition (0 h), we observed tdTomato-positive neurons in the DMH (48.8 ± 10.2 cells/mm^2^), VMH (16.9 ± 4.9 cells/mm^2^) and ARC (66.0 ± 13.0 cells/mm^2^) (Fig. 1c). There was no significant change in the number of tdTomato-positive DMH neurons when 4-OHT was injected 0.5 h after refeeding compared with the fasting only condition (0 h). However, the number of tdTomato-positive neurons in the DMH increased 2.88 and 2.22-fold when 4-OHT was injected 1 and 2 h after refeeding, respectively (Fig. 1d). There was no significant change in tdTomato-positive neurons in the VMH or ARC (Fig. 1d). The tdTomato-positive neurons in the PVH were localized to the dorsal region at 0 h, but were also detected in the parvocellular and magnocellular regions at 1 h after refeeding (Fig. 1c). Therefore, the distribution of tdTomato-positive neurons in the PVH were assessed according to the subdivisions described in the previous report^21^ (Fig. 1e). In the fasting only state (0 h), the number of tdTomato-positive neurons in the parvocellular and magnocellular regions were 209.2 ± 34.7 and 569.2 ± 59.3 cells/mm^2^, respectively. The number of tdTomato-positive neurons was increased 2.87-fold in the parvocellular region 1 h after refeeding, while no significant change was observed in the magnocellular region (Fig. 1f). Taken together, these results suggest that fasting–refeeding activates neurons in the DMH and the parvocellular region of the PVH.

### Chemogenetic activation of TRAP-labelled DMH neurons decreases food intake

To understand the roles of labelled DMH neurons responding to refeeding, we used the targeted expression of genetically modified DREADDs that are activated exclusively by clozapine (CLZ). Male Arg3.1-CreER^T2^ mice used in the excitatory DREADD study received bilateral Cre-inducible-G_q_-DREADD virus (AAV2-hSyn-DIO-hM3DGq-mCherry) or control virus (AAV2-hSyn-DIO-EGFP) injection into the DMH. After 2 weeks of recovery, animals were fasted overnight and re-fed RCD. 4-OHT was injected (i.p.) 1 h after refeeding to induce Cre-recombination (Fig. 2a). Experiments using CLZ were carried out 2 weeks after the 4-OHT injection to ensure the expression of the exogenous receptors.

Activation of neurons by CLZ injection was confirmed by immunohistochemistry (IHC) for cFos expression. After overnight fasting, the DMH of Arg3.1-CreER^T2^ mice injected with the excitatory DREADD or control virus (EGFP) received CLZ injection (i.p.) 1 h before perfusion. The percentage of colocalization of EGFP and cFos-expressing neurons was 4.17 ± 2.3% in the control group. The colocalization of excitatory DREADD and cFos-expressing neurons was 74.0 ± 2.5%, indicating that the DMH neurons expressing the excitatory DREADD were activated by CLZ stimulation (Fig. 2b, c).

The effect of DMH neuron activation on food intake was explored using CLZ stimulation. The Arg3.1-CreER^T2^ mice expressing the excitatory DREADD in the DMH were fasted overnight and treated with saline or CLZ 30 min before re-feeding with RCD. The food intake values for the saline-treated group at 1 and 2 h after refeeding were 1.08 ± 0.08 g and 1.30 ± 0.13 g, respectively. In the CLZ-treated group, the food intake decreased significantly at 1 and 2 h after refeeding compared with the saline-treated group (Fig. 2d). The effect of activated neurons on glucose metabolism was also evaluated using the glucose tolerance test (GTT) and the insulin tolerance test (ITT). The animals received CLZ or saline injection 30 min before the tests. The blood glucose levels among the groups remained similar during the GTT and ITT, indicating that the activation of these neurons has no impact on whole-body glucose metabolism. Our results show that the activation of refeeding-responsive DMH neurons reduces food intake without affecting glucose metabolism in mice.

### Chemogenetic inhibition of TRAP-labelled DMH neurons increases food intake

We further investigated the role of refeeding-responsive DMH neurons by using an inhibitory DREADD. Female Arg3.1-CreER^T2^ mice received bilateral injection of Cre-inducible G_i_-DREADD virus (AAV2-hSyn-DIO-hM4DGi-mCherry) into the DMH. After a 2-week recovery, the mice were fasted overnight and received 4-OHT injection (i.p.) 1 h after refeeding. One week after the 4-OHT injection, food intake was measured. After 16 h of fasting, animals received either saline or CLZ injection (i.p.) 30 min before being re-fed RCD. Food intake in the saline-treated mice after 1 and 2 h were 0.462 ± 0.038 g and 0.623 ± 0.060 g, respectively. The CLZ-treated mice significantly increased food intake at 1 and 2 h after refeeding (Fig. 3a). Saline and CLZ-treated groups had similar blood glucose levels in GTT and ITT (Fig. 3b, c), suggesting that the inhibition of DMH neurons responding to refeeding increased food intake, without affecting whole-body glucose metabolism.

**Figure 3.**
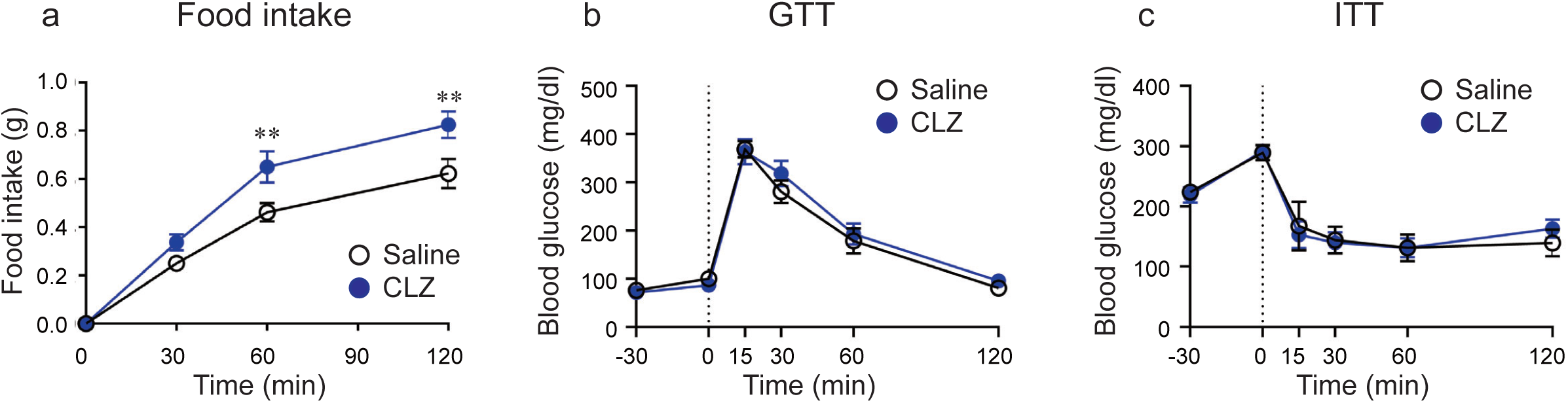
Chemogenetic inhibition of labelled DMH neurons increases food intake without impacting glucose metabolism. **a** Food intake (0–120 min) after i.p. injection (−30 min) of CLZ (*n* = 12) or saline (*n* = 13) in female Arg3.1-CreER^T2^ mice who received bilateral AAV2-hSyn-DIO-hM4DGi-mCherry virus injection into the DMH. 4-OHT was injected 1 h after refeeding. **b** GTT (0–120 min) after i.p. injection (−30 min) of CLZ (*n* = 6) or saline (*n* = 6). **c** ITT (0–120 min) after i.p. injection (−30 min) of CLZ (*n* = 6) or saline (*n* = 6). Each point represents mean ± SEM. Two-way ANOVA, Sidak post hoc test. \*\**P* < 0.01.

### Labelled DMH neurons project to the PVH

To identify the neural circuitry underlying the inhibition of food intake, we examined the projection site of the refeeding-responding DMH neurons using GFP-linked halorhodopsin (NpHR). NpHR is a light-driven chloride pump that is used as an inhibitory optogenetic actuator. NpHR is expressed in the cell body, as well as axons and their terminals, allowing identification of projection sites. Male Arg3.1-CreER^T2^ mice were injected Cre-inducible NpHR virus (AAV8-Syn-DIO-eNpHR3.0-EGFP) into the DMH (Fig. 4a). After a 2-week recovery period, the animals were fasted overnight and re-fed RCD. One hour later, 4-OHT was injected (i.p.). Two weeks later, animals were perfused for analysis. GFP expression in the DMH was detected, indicating that refeeding-responding neurons were successfully infected by the NpHR virus (Fig. 4b). Whole-brain regions were examined for GFP expression in the axon terminals. We observed GFP expression in the PVH, but not the ARC or other regions (Fig. 4c, d). This suggests that the direct projection from the DMH to the PVH regulates food intake.

**Figure 4.**
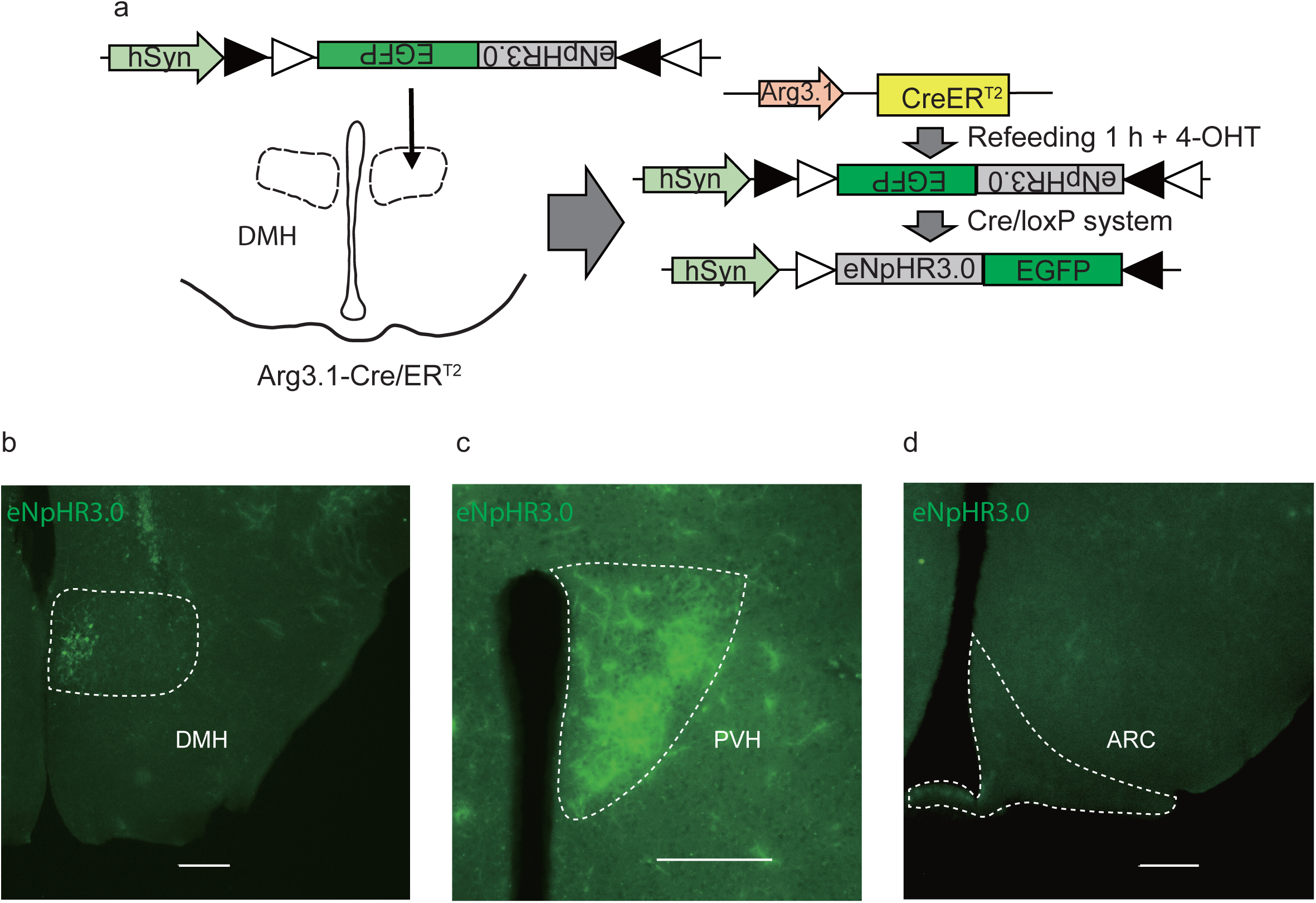
Refeeding-responsive DMH neurons project to the PVH. **a** The AAV vector construct used for this experiment. Male Arg3.1-CreER^T2^ mice received AAV8-Syn-DIO-eNpHR3.0-EGFP virus injection into the DMH. After recovery, the mice were fasted overnight and re-fed RCD. 1 h after refeeding, mice were given 4-OHT injection (i.p.) to induce Cre recombination (*n* = 3). **b** eNpHR3.0 (green) expression in the DMH (injection site). **c** eNpHR3.0 (green) expression in the PVH. **d** No expression of eNpHR3.0 (green) in the ARC.

### Chemogenetic activation of labelled DMH neurons induces conditioned place preference

Our results suggest that the labelled DMH neurons participate in the regulation of food intake. Previous studies indicate that the brain circuits responsible for controlling appetite are also involved in regulating emotional processes, including positive emotion triggered by reward ^22–24^. To investigate the role of labelled DMH neurons in emotion, we examined the effect of chemogenetic activation of these cells in the CPP test (Fig. 5a). Male Arg3.1-CreER^T2^ mice expressing excitatory DREADD (hM3DGq) or EGFP (control) in DMH neurons responsive to refeeding were used for this experiment. The CPP apparatus contained two chambers distinguished by the patterns on the floor (with or without stripes). During the tests, the mice were allowed to move freely between the chambers (Fig. 5a). On day 1 (pre-test), the mice were placed in the CPP apparatus for 10 min to determine intrinsic place preference. The time spent in each of the chambers was measured, and a heatmap was plotted. Before the conditioning phase, the PPIs for the two groups (hM3DGq: −5.8 ± 6.2%, *n* = 4; control: −3.9 ± 2.4%, *n* = 6 [mean ± SEM]) remained similar, indicating that both groups spent more time in the chamber with stripes (Fig. 5b, c).

**Figure 5.**
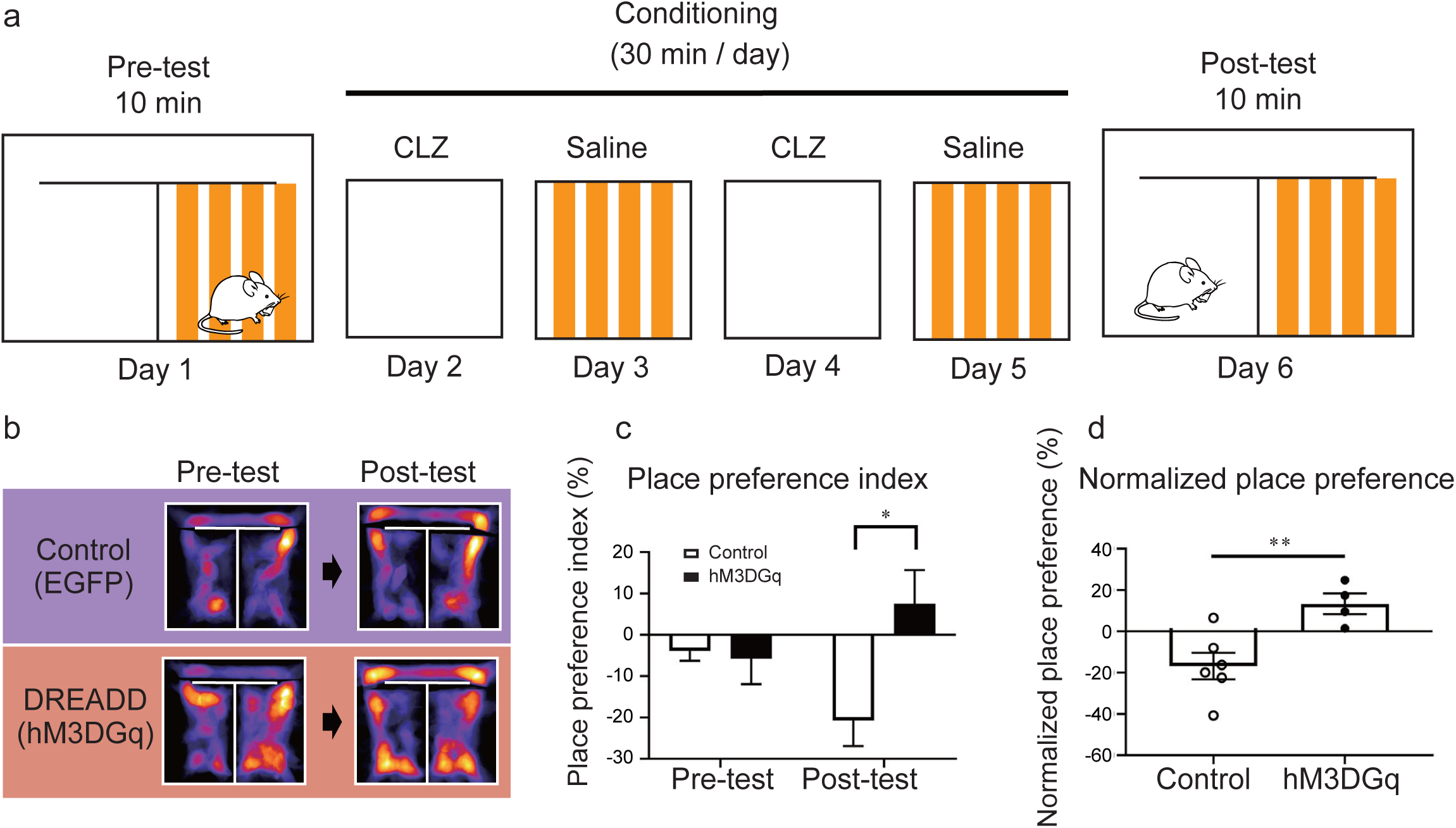
Intrinsic place preference is blocked by chemogenetic activation of labelled DMH neurons. **a** Scheme of the CPP test procedure: In the pre-test, excitatory DREADD (hM3DGq) or EGFP (control)-expressing male Arg3.1-CreER^T2^ mice were placed in the CPP apparatus for 10 min to determine intrinsic place preference (Day 1). The conditioning period took place on alternate days (Days 2–5). The mice were placed on one side of the chamber for 30 min and received either CLZ (nonpreferred side) or saline (preferred side) injection (i.p.) on each day. In the post-test, the mice were placed in the CPP apparatus for 10 min for place preference assessment (Day 6). **b** Representative heatmaps showing the location of EGFP (control) and excitatory DREADD (hM3DGq)-expressing mice in the pre- and post-tests. **c** Place preference index for EGFP (control) (*n* = 6) and hM3DGq (*n* = 4)-expressing mice. Data are expressed as mean ± SEM. Two-way ANOVA, Tukey post hoc test. \**P* < 0.05. **d** Normalized place preference for EGFP (control) (*n* = 6) and hM3DGq (*n* = 4)-expressing mice. Data are expressed as mean ± SEM. Unpaired *t*-test. \*\**P* < 0.01.

The conditioning took place on alternate days; on days 2 and 4, mice received CLZ injections (i.p.) and were placed in the nonpreferred side (without stripes) for 30 min with a barrier in the hallway. On days 3 and 5, mice received saline injections (i.p.) and were placed in the preferred side (with stripes) for 30 min. On day 6 (post-test), the mice were placed in the CPP apparatus for 10 min without the barrier, and place preference was measured (Fig. 5a). We observed a significant reduction in the post-test PPI in the control group compared with the hM3DGq group (Fig. 5c; *P* < 0.02), indicating the control group spent more time in the chamber with stripes. The normalized place preference shows that the hM3DGq group spent more time in the chambers without stripes in the post-test compared with the control group (Fig. 5d, *P* < 0.01). The conditioning phase had no impact on the preference in the control group, indicating that the development of CPP was not affected by CLZ injection. In the hM3DGq group, intrinsic place preference was blocked by chemogenetic activation of DMH neurons, resulting in more time spent in the previously nonpreferred side (without stripes) (Fig. 5b– d). Taken together, our results suggest that refeeding-responsive DMH neurons are involved in encoding the positive valence induced by refeeding.

### Pdyn neurons in the DMH are responsive to refeeding

The hypothalamus comprises various cell types, and their transcriptional profiles provide key information towards understanding this region’s function in homeostatic regulation^19^. To investigate the transcriptional profile of tdTomato-positive DMH neurons, we performed an RNA-sequencing study using cells dissociated from male Arg3.1-Cre/ER^T2^;Ai14 mice.

After overnight fasting, mice were fed RCD and given 4-OHT injection (i.p.) 1 h after refeeding. The tissue containing the DMH was dissected, and after cell dissociation, tdTomato-positive and negative (control) DMH neurons were sorted manually (5 cells per sample) for sequencing (Fig. 6a). We generated gene expression heatmaps based on previous findings^19^. Pan neuronal markers, including Snap25 and Syt1, were detected in all samples (tdTomato-positive, *n* = 4; control, *n* = 4) (Fig. 6b), suggesting successful sorting of tdTomato-negative (control) neurons, thereby permitting assessment of differential transcripts. As the probability of tdTomato-negative neurons being other non-neuronal cell types is high, relatively low expression of non-neuronal marker genes was observed in the control samples (Fig. 6b). The expression of the glutamatergic neuron marker Slc17a6 was higher than that of the GABAergic neuron marker Slc32a1 in tdTomato-positive neurons (Fig. 6b), suggesting that the refeeding-responding DMH neurons are glutamatergic. We also assessed the expression profile of neuropeptides and receptors, and found high expression of CCK in tdTomato-positive neurons (Fig. 6c). According to a previous study^19^, glutamatergic neurons in the hypothalamus can be divided into 15 subtypes, and one of the markers is CCK. CCK-positive neurons express other markers (shown in Fig. 6d). However, the gene profile of tdTomato-positive neurons did not match those of the subtypes listed in that study^19^, indicating that either each sample contained neurons from multiple subtypes or that the labelled DMH neurons were an unidentified subtype.

**Figure 6.**
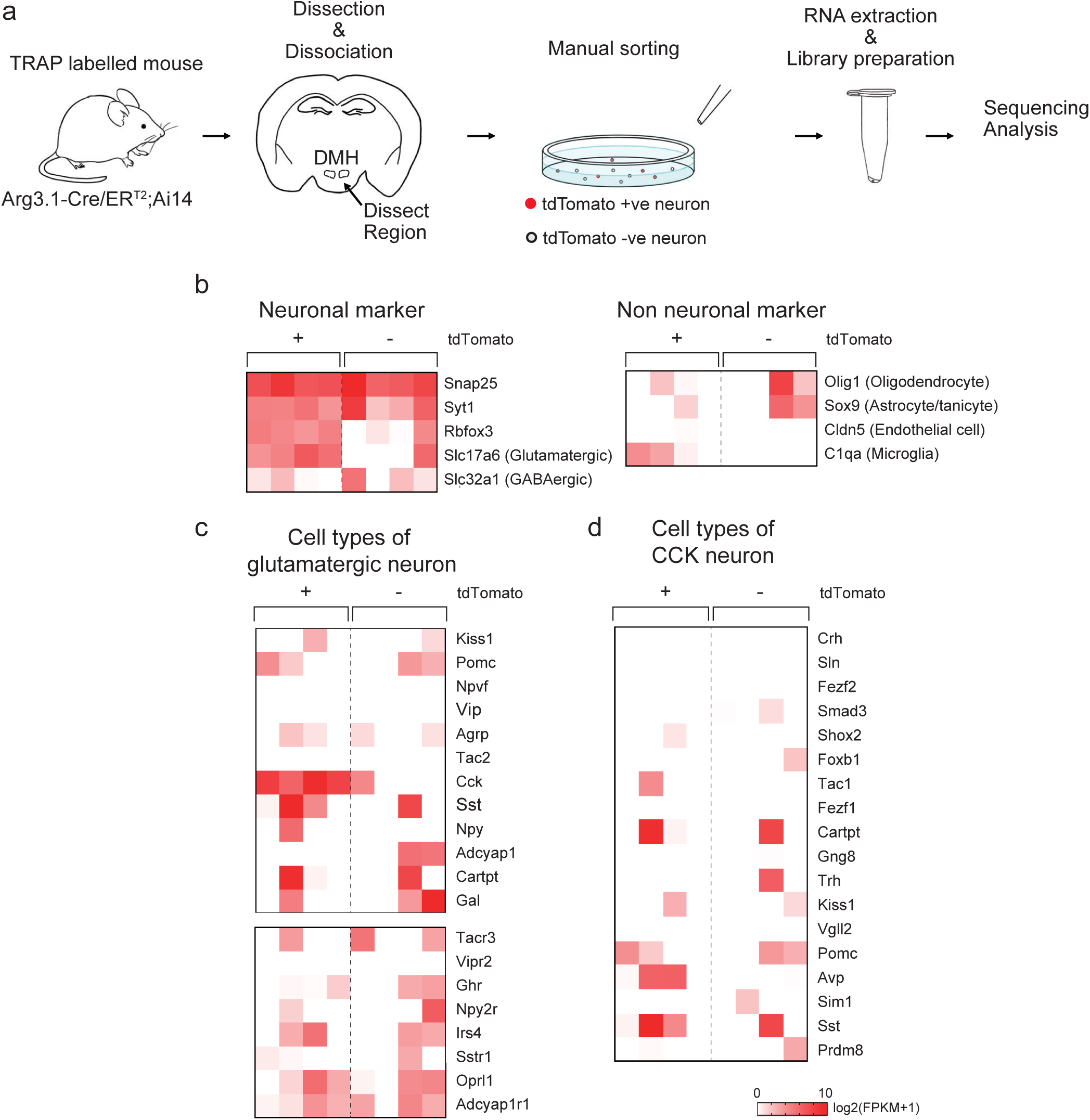
Refeeding-responsive DMH neurons express glutamatergic neuron markers. **a** Experimental scheme of the RNA sequencing protocol. Arg3.1-Cre/ERT2;Ai14 mice were TRAPed 1 h after refeeding. The DMH region was dissected and triturated for cell sorting. tdTomato-positive (+ve) and negative (−ve) neurons were collected manually for RNA sequencing. **b** Heatmaps representing the expression of neuronal and non-neuronal marker genes in tdTomato-positive and negative neurons. Log_2_ (FPKM + 1). **c, d** Heatmaps of the genetic markers for hypothalamic glutamatergic neurons. The gene list was obtained from a previous report^19^. Log_2_ (FPKM + 1).

To further investigate the unique transcriptional profile of tdTomato-positive neurons, we examined the gene expression of known ligands and receptors playing roles in cell–cell communication^20^. The gene expression heatmap for ligands were generated, and high expression of pdyn, gastrin-releasing peptide (GRP) and CCK were identified in tdTomato-positive neurons (Fig. 7a). Pdyn is a precursor of dynorphins, the endogenous ligands for opioid receptors. Immunohistochemistry showed that 68.6 ± 9.6% (mean ± SEM, *n* = 3) of tdTomato-positive neurons colocalized with dynA (Fig. 7d).

**Figure 7.**
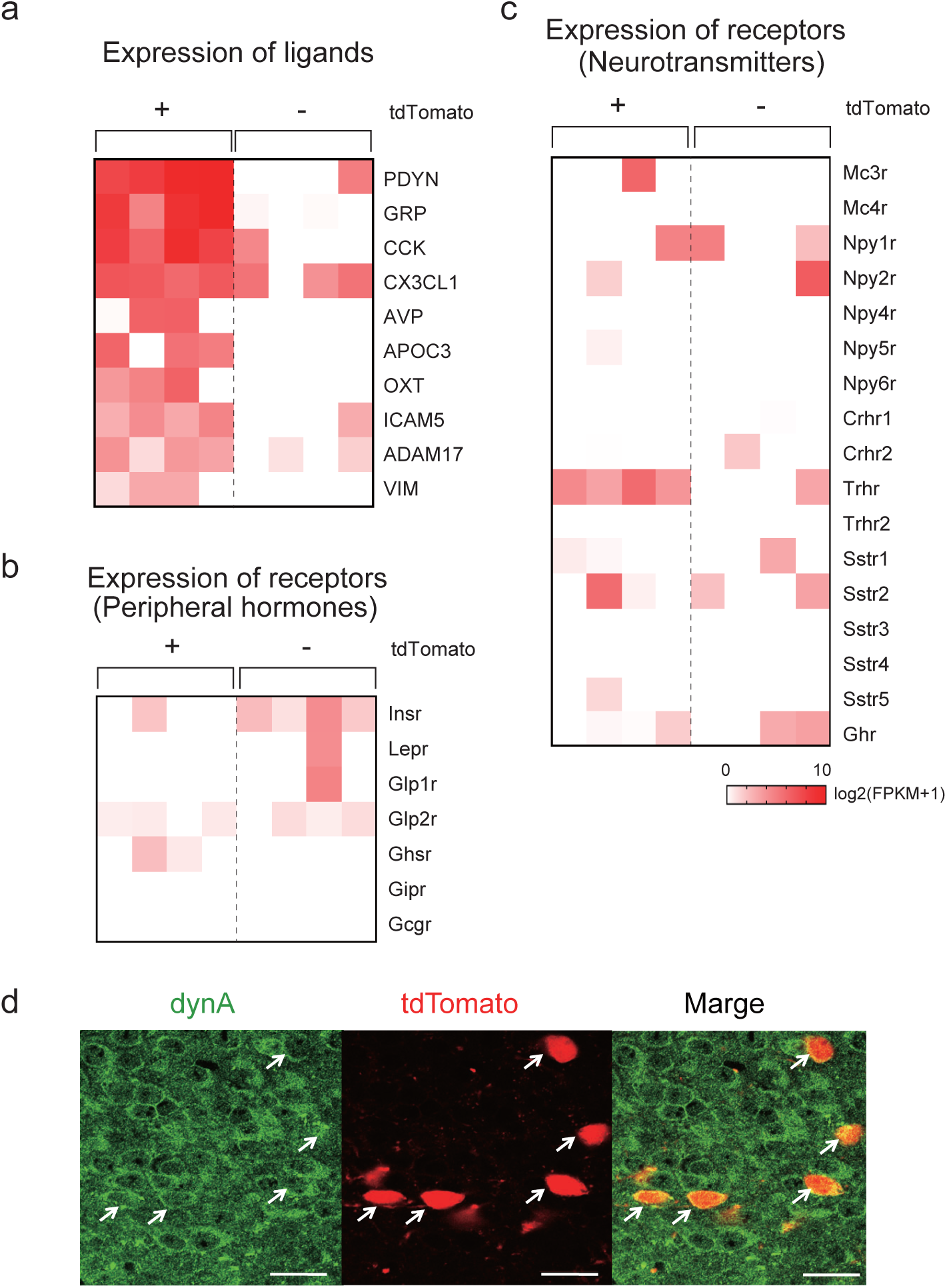
Refeeding-responsive DMH neurons express pdyn and CCK. **a** Heatmap of the gene expression of ligands and receptors involved in cell–cell communication. The gene list was obtained from a previous report ^20^. Log_2_ (FPKM + 1). **b, c** Gene expression of receptors for peripheral hormones (b) and neurotransmitters (c) involved in metabolism. **d** Representative micrographs showing dynA (green) colocalization with TRAP-labelled tdTomato (red)-expressing neurons in the DMH. Scale bar: 20 μm.

We also examined the expression of receptors for peripheral hormones and neurotransmitters (Fig. 7b, c), and detected expression of the thyrotropin-releasing hormone receptor (Trhr) gene in tdTomato-positive neurons. The expression of receptors for peripheral metabolic hormones, including the insulin (Insr), leptin (Lepr) and glucagon-like peptide-1 (Glp1r) receptors, were not detected (Fig. 7b). Collectively, our sequencing results suggest that refeeding-responsive DMH neurons labelled by the TRAP method are glutamatergic neurons expressing pdyn, GRP, CCK and Trhr.

## Discussion

Here, we identified a population of neurons in the compact part of the DMH activated by refeeding using the TRAP method. Chemogenetic activation and inhibition of TRAP-labelled neurons in the DMH decreased and increased food intake, respectively. The activation of the DMH neurons also promoted positive valence. The DMH neurons were identified as glutamatergic neurons that project to the PVH and express the genes for pdyn, GRP, CCK and Trhr. Our findings show that these novel DMH neurons have an important role in regulating food intake and are possibly involved in the satiety process for appetite and affectivity.

Homeostatic control of appetite is a complex process in which signals from peripheral hormones and the CNS are integrated to regulate feeding behaviours. Previous studies have shown that various subregions of the hypothalamus, including the ARC, VMH, PVH and lateral hypothalamus (LH), play important roles in the regulation of food intake. For example, POMC^8,9^, NPY/AgRP^25,26^ and dopamine neurons^27^ in the ARC, steroidogenic factor 1 (SF1) neurons^28^ in the VMH, corticotropin-releasing hormone neurons^16,29,30^ in the PVH, and orexin neurons^31^ in the LH have been reported to play roles in the regulation of feeding. Among these, POMC and NPY/AgRP neurons are considered first-order neurons that control feeding behaviour, because genetic deletion of hormone receptors in these neurons significantly impacts feeding behaviour and body weight. However, these neurons quickly respond to the onset of feeding or even the perception of food^32,33^. Therefore, it is unclear which neurons generate satiety.

The TRAP method allows one to examine neurons based on activity induced by refeeding. We used Arg3.1-CreER^T2^ because Arg3.1 has never been used in the study of the hypothalamus, and it could therefore identify novel cell populations. Although activation of SF1 and POMC neurons can terminate feeding behaviour^5,28^, TRAP-labelled neurons in the VMH and ARC displayed a similar activation status in both fasted and re-fed states in the present study. While some POMC and VMH neurons have been reported to express Arg3.1 in a model of inflammation^34^, these neurons were not activated by refeeding. However, POMC and VMH neurons have been reported to express cFos after refeeding^35^. Thus, the TRAP-labelled DMH neurons likely regulate satiety together with neurons in other hypothalamic regions, such as the ARC and VMH.

The gene expression profile of TRAP-labelled DMH neurons resembled that of CCK neurons reported in a previous study^19^. However, our RNA sequencing analysis suggests their gene expression profile is nonetheless distinct from that of the reported subtypes. The secretion of the peripheral hormones insulin and GLP1 increases after food intake, and they modulate neural function to decrease hunger and food intake^36,37^. However, expression of the receptors Insr, Lepr and Glp1r was not observed, suggesting that DMH neurons receive other afferent signals to regulate food intake (Fig. 7). Trhr is the only receptor that was expressed specifically in the TRAP-labelled neurons. A subset of TRH neurons in the PVH is activated by refeeding^38^, and injection of TRH into the third ventricle or medial hypothalamus, including the DMH, suppresses food intake after fasting^39^. Therefore, TRH neurons may be involved in regulating the activity of TRAP-labelled neurons in the DMH.

Pdyn is a precursor of dynorphins, which are endogenous opioid peptides that signal via mu (MOP), kappa (KOP) and delta opioid (DOP) receptors. Dynorphins have a high affinity for KOP receptors, and activation of KOP receptors has an antinociceptive effect^40^. Dynorphins are also associated with negative valence, as KOP receptor activation in the nucleus accumbens (NAC) decreases dopamine release^41,42^. Previous studies suggest that the central opioid system is involved in the regulation of feeding behaviour^43^, and have highlighted the role of MOPs and other opioids, enkephalin and beta-endorphins, but not KOPs or dynorphins, in food intake and the rewarding effects of food^43,44^.

Our current findings demonstrate that the activation of TRAP-labelled pdyn/CCK neurons in the DMH promotes positive valence and inhibits food intake. This suggests that dynorphins released by DMH neurons bind to MOP and/or DOP receptors, but not KOP receptors. MOP and DOP receptors are involved in the rewarding effect of the opioid system. Although dynorphins have a high affinity for KOP receptors, they can also interact with other opioid receptors^40^. Further investigation is necessary to identify the opioid receptors involved in the positive affect induced by the activation of DMH neurons.

Consistent with our findings, previous reports have implicated pdyn neurons in energy homeostasis. A study by Allison et. al. identified neurons in the DMH expressing Lepr/pdyn using translating ribosome affinity purification and RNA sequencing analysis^45^. Ablation of Lepr in pdyn neurons alters energy expenditure in mice^45^. Garfield et. al. demonstrated that Lepr/pdyn-expressing GABAergic DMH neurons project to AgRP neurons in the ARC and modulate feeding behaviour^46^. However, in the current study, we did not detect Lepr expression or projections to the ARC, indicating that pdyn neurons labelled by the TRAP method are a different subpopulation of neuron. A study using ribosome phosphorylation identified pdyn-expressing neurons in the DMH that were activated during scheduled feeding^47^. Intracerebroventricular injection of a KOP receptor antagonist, norbinaltorphimine, increased food intake during scheduled feeding^47^. Taken together, these observations suggest that the DMH contains several subtypes of pdyn neurons that are involved in the regulation of feeding behaviour and energy expenditure.

There are two forms of appetite, homeostatic and hedonic appetite, and their regulatory mechanisms affect each other to maintain the balance between food intake and energy expenditure^48^. The experiencing of pleasure is a key factor in determining future behavioural actions^43^. Our findings indicate that activation of pdyn/CCK DMH neurons inhibits food intake and also promotes positive valance, suggesting a key role of emotion in the transition from hunger (desire to eat) to satiety (termination of hunger). Numerous studies have reported on the rewarding effect of food and the role of mesolimbic pathways in producing the hedonic response^49^. Our finding suggests that pdyn/CCK neurons in the DMH play an important role in connecting the feeling of satisfaction and the termination of feeding behaviour. To further elucidate the role of emotion, additional study of the neuronal networks upstream and downstream of the refeeding-responsive DMH neurons is needed.

In summary, we identified refeeding-responding pdyn/CCK neurons in the DMH using the TRAP method. Activation of these neurons decreased food intake, implicating them in satiety. Our findings suggest that the hypothalamic opioid system is a potential novel therapeutic target for obesity.

## Acknowledgments

We thank Takako Usuda for providing the mouse illustration. This work was supported by Leading Initiative for Excellent Young Researchers (from MEXT); a Grant-in-Aid for Young Scientists (A) (Grant Number JP17H05059), a Grant-in-Aid for Scientific Research (B) (Grant Number JP18H02857); Japanese Initiative for Progress of Research on Infectious Diseases for Global Epidemics (JP17fm0208011h0001, JP18fm0208011h0002, JP19fm0208011h0003); the Takeda Science Foundation; the Uehara Memorial Foundation; Astellas Foundation for Research on Metabolic Disorders; Suzuken Memorial Foundation and Program for supporting introduction of the new sharing system (JPMXS0420100617, JPMXS0420100618, JPMXS0420100619).

## Author Contributions

C.T. conceived this study and designed the experiments. D.I. performed most of the experiments, and C.T. supervised the entire study. I.Y. and H.M. performed the TRAP, GTT and ITT experiments. D.I. and T. Yonekura injected AAVs. D.I. and K. Kato conducted the CPP. I.Y., T. Yamasaki, T. Yonekura, K.O. and M.H. performed parts of the single-cell RNA sequence study. I.Y., D.I. and C.T. performed data analysis. I.Y., D.I., N.I., K. Kimura and C.T. wrote or contributed to the writing of the manuscript.

## Materials & Correspondence

Further information and requests for resources and reagents should be directed to and will be fulfilled by the lead contact, Chitoku Toda.

